# A Unified Framework for Variance Component Estimation with Summary Statistics in Genome-wide Association Studies

**DOI:** 10.1101/042846

**Authors:** Xiang Zhou

## Abstract

Linear mixed models (LMMs) are among the most commonly used tools for genetic association studies. However, the standard method for estimating variance components in LMMs – the restricted maximum likelihood estimation method (REML) – suffers from several important drawbacks: REML is computationally slow, requires individual-level genotypes and phenotypes, and produces biased estimates in case control studies. To remedy these drawbacks, we present an alternative framework for variance component estimation, which we refer to as MQS. MQS is based on the method of moments (MoM) and the minimal norm quadratic unbiased estimation (MINQUE) criteria, and brings two seemingly unrelated methods – the renowned Haseman-Elston (HE) regression and the recent LD score regression (LDSC) – into the same unified framework. With this new framework, we provide an alternative but mathematically equivalent form of HE that allows for the use of summary statistics and is faster to compute. We also provide an exact estimation form of LDSC to yield unbiased and more accurate estimates with calibrated confidence intervals. A key feature of our method is that it can effectively use a small random subset of individuals for computation while still producing estimates that are almost as accurate as if the full data were used. As a result, our method produces unbiased and accurate estimates with calibrated standard errors, while it is computationally efficient for large data sets. Using simulations and applications to 33 phenotypes from 7 real data sets, we illustrate the benefits of our method for estimating and partitioning chip heritability. Our method is implemented in the GEMMA software package, freely available at www.xzlab.org/software.html.

## Introduction

Linear mixed models (LMMs), sometimes referred to as variance component models, have been widely applied in many areas of genetics. For example, they have been used for linkage analysis and heritability estimations in family studies [1–5], for association analysis to control for individual relatedness and population stratification [6–15], for genomic selection and risk prediction by jointly modeling genome-wide SNPs [16–22], and for rare variant association tests by grouping individually weak effects to improve power [23]. More recently, with growing interest, LMMs have been applied to estimate the proportion of phenotypic variance explained by available SNPs [21, 22, 24–26] – a quantity often referred to as chip heritability – and to partition the chip heritability by different chromosome segments or by different functional genomic annotations [27–30]. These applications all require accurate estimation of variance components in LMMs. However, due to the increasingly large samples being collected today, estimating variance components in genetic data is becoming increasingly challenging.

Two standard statistical methods exist for variance component estimation. The first is the restricted maximum likelihood estimation (REML) method. REML is statistically efficient, but has three major drawbacks. First, despite recent computational innovations [7, 10–12, 31], REML is computationally expensive. It requires not only an iterative optimization algorithm that scales cubically with the number of individuals, but also a pre-computation step to construct a genetic relatedness matrix from genome-wide SNPs – a step that scales both quadratically with the sample size and linearly with the number of markers. Second, REML requires individual-level genotypes and phenotypes, both of which are not readily available in many large consortium studies or meta-analyses. Using individual-level data thus restricts the use of REML and limits the benefits of LMMs in large-scale studies. Finally, REML relies on the normality assumption of residual errors and is not robust to model misspecification. In particular, in ascertained case control studies or studies with an extreme sample design, REML underestimates chip heritability [32,33].

A long existing alternative to REML for variance component estimation is the minimal norm quadratic unbiased estimation (MINQUE) method, a method of moments (MoM) [34,35]. Because the MINQUE estimates are relatively efficient and closely related to REML estimates [36,37], MINQUE is widely used in many non-genetic settings and has also been applied to animal breeding programs [38]. However, MINQUE shares the drawbacks of REML: it requires computing a genetic relatedness matrix and is thus computationally slow, it does not allow the use of summary statistics, and it has unknown properties in case control studies.

To remedy these three drawbacks, we present a new, alternative framework for variance component estimation. We refer to our method as MQS (MinQue for Summary statistics) as it is based on MINQUE, but addresses its shortcomings with two approximations. Our first approximation allows us to use summary statistics and obtain unbiased estimates at the price of a small reduction of estimation accuracy. Our second approximation allows us to use only a small random subset of individuals to compute the genetic relatedness matrices. This sub-sampling strategy greatly improves computational efficiency, but does not significantly reduce the estimation accuracy in MQS. Most importantly, our framework unifies two seemingly unrelated methods – the renowned Haseman-Elston (HE) regression [32,39-44] and the recent LD score regression (LDSC) [30,45,46] – into the same umbrella. With simulations and applications to 33 phenotypes from 7 real data sets, we illustrate the benefits of our method.

## Results

### Method Overview

Our method applies to the following LMMs that can be used to partition chip heritability into *k* different non-overlapping categories

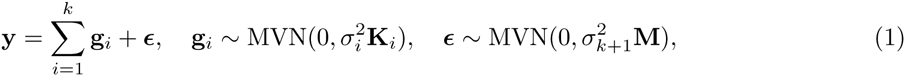

where **y** is an *n*-vector of phenotypes for *n* individuals; **g**_*i*_ is an *n*-vector of random effects representing the combined genetic effects of SNPs in *i*th category; 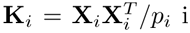 is an *n* by *n* genetic relatedness matrix computed from the *n* by *p*_*i*_ genotype matrix for *p*_*i*_ SNPs in ith category; 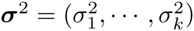 are the corresponding variance components; *ϵ* is a *n*-vector of residual errors; 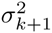 is the residual error variance; 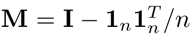 is a projection matrix; and MVN denotes a multivariate normal distribution. Both **y** and every column of **X** have been centered to have mean zero, allowing us to ignore the intercept and use **M** instead of the usual identity matrix **I** to constrain the errors to have mean zero. We denote the phenotype variance 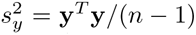. We also define the scaled version of the variance components as 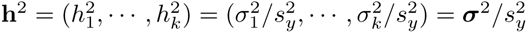.

We have described our method for variance component estimation in detail in the Methods section. Briefly, our framework, which we refer to as MQS, is based on MINQUE and provides unbiased estimates with calibrated standard errors. MQS has a simple, closed-form solution for estimating the *k* variance components: **σ̂**^2^ = **S**^‒1^**q**, with a *k*-vector **q** and a *k* by *k* matrix **S** (equation 11). Intuitively, the ith element of **q** measures the proportion of variance in phenotypes explained (PVE) by SNPs in *i*th category when SNPs are independent, while *ij*th element of **S** accounts for the linkage disequilibrium (LD) between SNPs in *i*th and *j*th categories. MQS requires the marginal z-scores for computing **q**, the usual genetic relatedness matrices for computing **S** (or the genetic relatedness matrices from a reference panel for estimating **S**; see below), and a set of pre-specified SNP weights that are used for both **q** and **S**. This set of pre-specified SNP weights is used in MQS to approximate the optimal MINQUE estimating equations. Different choices of weights represent different ways of approximation and can lead to unbiased estimates with different accuracy.

MQS is a unified framework because different SNP weighting options lead to different estimation methods. We consider two particular weighting options here. The first option is equal SNP weights (equation 9). We refer to the variation of MQS under this weighting option MQS-HEW. MQS-HEW is mathematically equivalent to the renowned Haseman-Elston (HE) cross-product regression [32,39–44]. However, our particular MQS formulation allows us to both make use of summary statistics and compute the standard errors faster than before. The equivalence between HE and MQS-HEW guarantees the MQS-HEW estimates to be unbiased for case control studies [32,33]. The second option assigns SNP weights as a function of both the LD scores and *a priori* set of variance components (equation 10). We refer to the variation of MQS under this weighting option MQS-LDW. For *k* = 1, MQS-LDW is effectively an exact form of LDSC [45]. However, in contrast to LDSC, MQS-LDW uses **S** instead of LD scores to measure SNP correlations. Because LD scores are inevitably under-estimated, using LD scores to measure SNP correlations is expected to yield biased estimates. In particular, when *k* = 1, LDSC will over-estimate the variance components. When *k* > 1, the variance components from LDSC will show different degrees of bias. By providing an exact form of LDSC and a new computing form of standard errors (CI1; see below), MQS-LDW yields unbiased and more accurate estimates with calibrated confidence intervals.

One particular feature of MQS is that we can use a random subset of individuals to obtain an estimate of **S** – **Ŝ** – without significantly reducing the accuracy of the variance component estimates. Estimating **S** with a smaller sub-sample instead of computing **S** with the full data not only improves computational efficiency, but also allows us to apply MQS to data from many consortium studies: thus, we can pair **q** computed from the available marginal z-scores in the consortium study with **Ŝ** estimated from a random sub-sample of the study. When such a random sub-sample of the study is not available, we can also use a reference panel, such as the 1000 genome project [47], to estimate **S**, so long as individuals in the reference panel can be viewed as a sub-sample of the study (e.g. of the same ethnic origin). Our statistical arguments for using the sub-sampling strategy, as detailed in the Methods section, is based on examining the variances of **q** and **Ŝ**. Briefly, the variance of **σ̂**^2^, ***V***(**σ̂**^2^), is contributed by both ***V***(**q**) and***V***(**Ŝ**). However, ***V***(**q**) dominates the contribution and can be 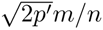 times larger than ***V***(**Ŝ**), where *p*′ is the effective number of independent SNPs and *m* is the number of sub-samples. Thus, we can use a much smaller number of individuals *m* to estimate **S** without increasing ***V***(**σ̂**^2^) significantly. In addition, our computation of standard errors explicitly accounts for the uncertainty introduced by using a much smaller set of individuals to estimate **S** instead of computing it.

Finally, MQS provides three different options to compute the standard errors. The first option, CI, requires genetic relatedness matrices from the full data but is *n* times faster than the standard formula used in MoM (equations 15, 16 and 18). The second (CI1) and the third (CI2) options use only summary statistics (equations 20 and 21). CI1 is exact but requires additional summary statistics besides the marginal z-scores equation (equation 25). CI2 follows the approach of LDSC [45] and requires only permutation of the marginal z-scores. However, CI2 assumes SNP independence and is approximate. Thus, CI2, like LDSC, does not yield calibrated confidence intervals when the LD pattern is complicated. Examples where CI2 does not apply include ascertained case control studies [48,49], admixture populations [50], and related individuals (defined loosely as individuals who are not far away in time from their most recent common ancestor, MRCA [51]).

We summarize the key features of several variance component estimation methods and their computational complexity in Table 1.

**Table 1.**
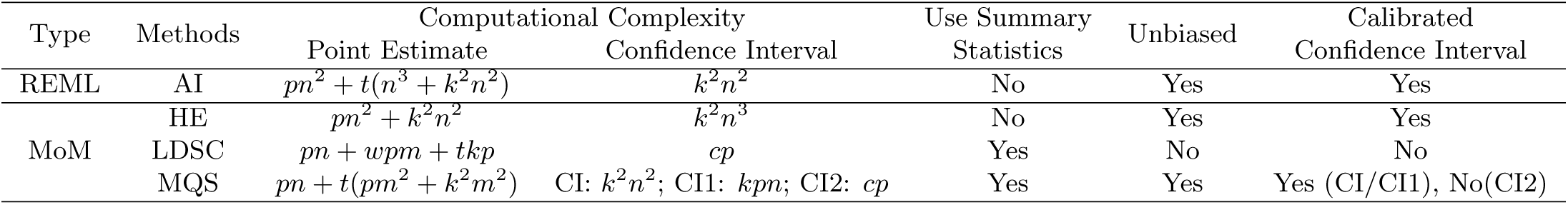
Features and computational complexity of different methods for variance component estimation. The computational complexity includes time to compute the genetic relatedness matrices. *n* is the number of individuals, *m* is the number of randomly selected subset of individuals (*m* < *n*), *k* is the number of variance components, *p* is the number of genetic markers. AI: average information method. HE: Haseman-Elston regression. LDSC: LD score regression. *w* is the average number of variants used to estimate the LD scores. *c* is the number of blocks used for the block-wise Jackknife re-sampling algorithm. t is the number of iterations used for estimation: *t* = 1 for MQS-HEW, *t* = 2 (or *t* > 2) for MQS-LDW and LDSC. The confidence interval in MQS can be computed in three different ways: CI requires computed genetic relatedness matrices; CI1 requires summary statistics in addition to z-scores; CI2 follows LDSC, requires only z-scores but is an approximation.

### Simulations: Point Estimates and Confidence Intervals

We perform simulations to compare the performance of several different methods on variance component estimation. We use two real genotype data sets for simulations: an Australian data set with *n* = 3, 925 individuals and *p* = 4, 352, 968 imputed SNPs [24], and a Finish data set with *n* = 5,123 individuals and *p* = 319,148 genotyped SNPs [52]. We choose these two data sets not only because both consist of Caucasian individuals, but also because the two differ in LD pattern: the Finland data displays longer LD than the Australia data (Figure S4). The pattern of long LD in the Finland data makes it easy to validate some of our expectations. The Finland data displays such long LD pattern presumably because individuals from the Finland data are more closely related to each other than individuals from the Australia data (notice that the Finland study is not a family study however). For each data set, the real genotypes are used to compute genetic relatedness matrices, with which we simulate phenotypes based on LMMs.

We compare five different methods: (1) REML uses individual-level phenotypes and genotypes; (2) HE regression uses individual-level phenotypes and genotypes; (3) LDSC uses z-scores computed from the full data and LD scores estimated based on 1 MB window from the full data; (4) MQS-HEW uses z-scores computed from the full data, and S estimated from *m* = 400 randomly selected individuals; and (5) MQS-LDW uses z-scores computed from the full data, LD scores estimated based on 1 MB window from genotypes of *m* = 400 randomly selected individuals, and **Ŝ** estimated from the same *m* = 400 selected individuals. Notice that for MQS-HEW and MQS-LDW, a different set of *m* = 400 individuals are used in each replicate.

We first simulate phenotypes under LMMs with *k* =1. We check three scenarios: *h*^2^ = 0, 0.25, or 0.5 (notice that most quantitative traits have a chip heritability below 0.5 [53]). For each scenario, we perform 1,000 replicates. We obtain the variance component estimates from different methods. Figures 1A and 1C show boxplots of these estimates. Figures 1B and 1D show the accuracy of these estimates by plotting the mean squared error (MSE) relative to REML. The simulations validate a few of our expectations.

**Figure 1.**
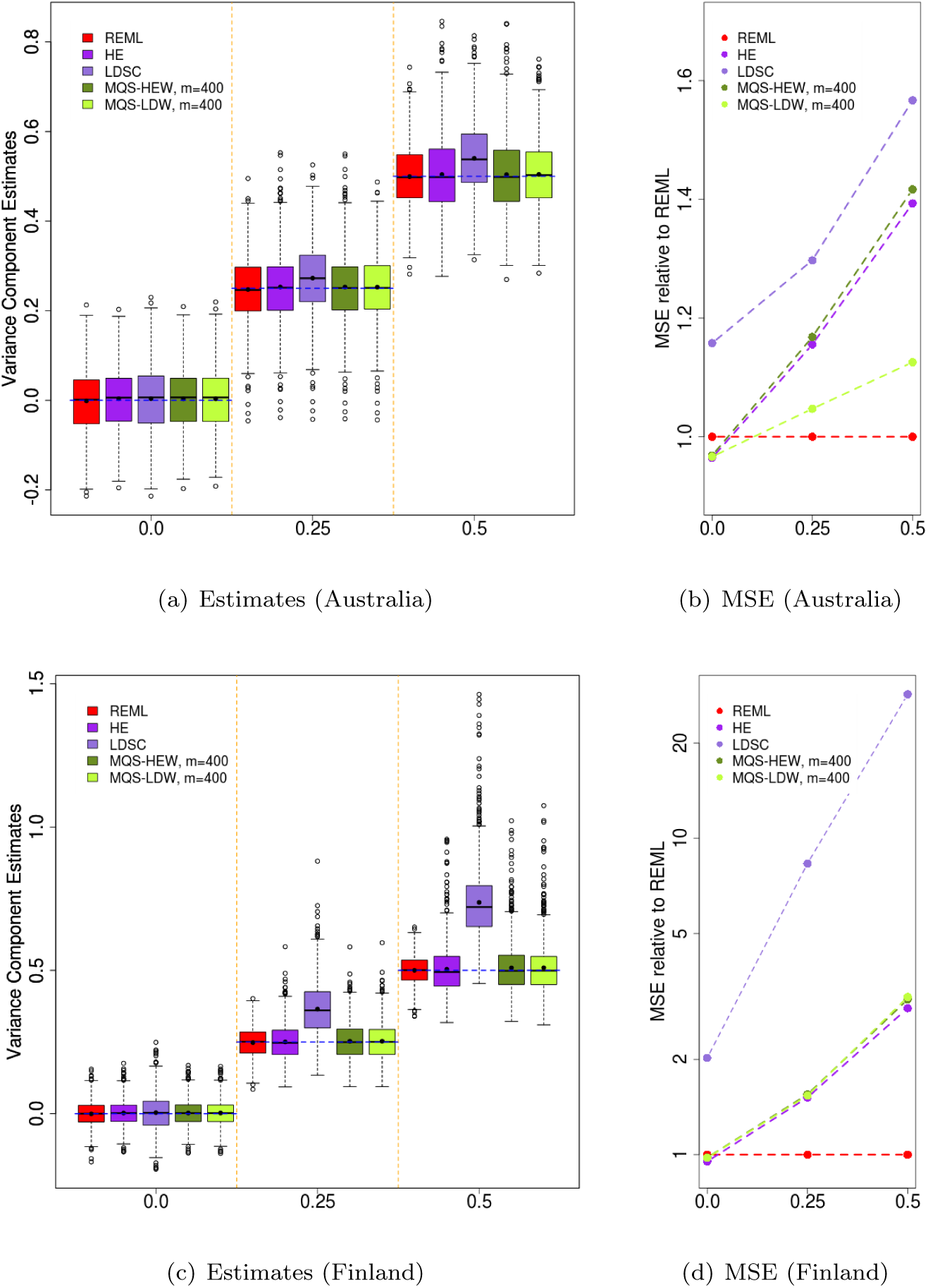
Comparison of variance component estimates from REML (red), HE (purple), LDSC (light purple), MQS-HEW (green), and MQS-LDW (light green) for *k* = 1 simulations based on the Australian data (a, b) or the Finland data (c, d). (a) and (c) show boxplots while (b) and (d) show the MSE relative to REML. The true variance components from the three simulations (0, 0.25 and 0.5) are shown on the x-axis of all panels, and are also shown as three blue horizontal dashed lines in (a) and (c). The dot in the middle of the boxplots represents the mean of the estimates in (a) and (c). Note that the y-axis in (d) is on log scale.

First, because MQS-HEW with **S** is identical to HE and because **Ŝ** is an accurate estimate of **S**, estimates from MQS-HEW with **Ŝ** are similar to those from HE. In addition, because the variance of the MQS estimates depends on a product of ***V***(**Ŝ**) and heritability, estimates from MQS-HEW are more similar to HE for a small *h*^2^ than for a large *h*^2^ (Figure S5).

Second, in practice LD scores are estimated via a sliding window based approach and are inevitably under-estimated. Because LDSC uses these under-estimated LD scores to approximate **S**, LDSC underestimates **S** and over-estimates the variance components. Such upward bias is more obvious in the Finland data than in the Australia data because of longer LD in the Finland data. For instance, when *h*^2^ = 0.5, the LDSC estimates on average are 8.0% higher than expected in the Australia data and 47.4% higher in the Finland data. Estimates from all other methods are unbiased. Because of the bias, the estimates from LDSC are the least accurate ones in all scenarios.

Third, MQS-HEW/HE and MQS-LDW approximate MINQUE in different ways, but both approximations are only accurate when the variance component is small and/or when individuals are unrelated. Thus, both MQS-HEW/HE and MQS-LDW are more accurate for a small *h*^2^ than for a large *h*^2^, and more accurate in the Australia data than in the Finland data. In addition, in the case of *h*^2^ = 0, both MQS-HEW/HE and MQS-LDW are more accurate than REML because they effectively assume *h*^2^ = 0 *a priori* and are locally optimal.

Fourth, presumably because MQS-LDW uses estimates from MQS-HEW as initial values, and presumably because MQS-LDW uses the extra information of LD scores to compute the SNP weights, MQS-LDW is more accurate than MQS-HEW in the Australia data. However, because long LD reduces the accuracy of LD score estimates, MQS-LDW is of similar accuracy as MQS-HEW in the Finland data.

Next, we simulate phenotypes under LMMs with *k* = 6. We first annotate the genome using six categories as in [29]. The six categories include coding, untranslated region (UTR), promoter, DNAse hyper-sensitivity regions (DHS), intronic and intergenic regions. SNPs are classified into these six categories based on genomic location. We then compute a genetic relatedness matrix for each category, based on which we simulate phenotypes using LMMs with multiple variance components. We examine three different scenarios: (I) a null scenario where all variance components are zero (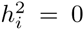 for *i* = 1,…,6); (II) an alternative scenario with total chip heritability 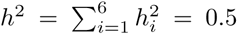, and with each category explaining an equal proportion (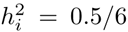 for *i* = 1,…, 6); (III) a more realistic alternative scenario with *h*^2^ = 0.5 and with DHS explaining a large proportion of heritability 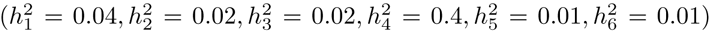. For each scenario, we perform 1,000 replicates. Figure 2 show the estimates and estimation accuracy for scenario III. Figures S6 and S7 show the estimates and estimation accuracy for scenario I and scenario II, respectively. The conclusions from these *k* = 6 simulations are largely consistent with *k* = 1 simulations. For example, all estimates except for LDSC are unbiased. Estimates from LDSC for the total heritability are upward biased. In particular, in scenario III, the total chip heritabilty estimates on average are 6.0% higher than expected in the Australia data and 45.9% higher than expected in the Finland data. However, *k* = 6 simulations also revealed new insights. First, because LD scores in different categories are estimated with different accuracy, individual variance component estimates from LDSC sometimes are upward biased and sometimes downward biased. Second, MQS-LDW is more accurate than MQS-HEW in scenario II with the Australia data, is worse than MQS-HEW in scenario I with the Australia data, and is of comparable accuracy as MQS-HEW in other cases. The comparison between MQS-LDW and MQS-HEW suggests that MQS-LDW is not always more accurate than MQS-HEW even in the Australia data.

**Figure 2.**
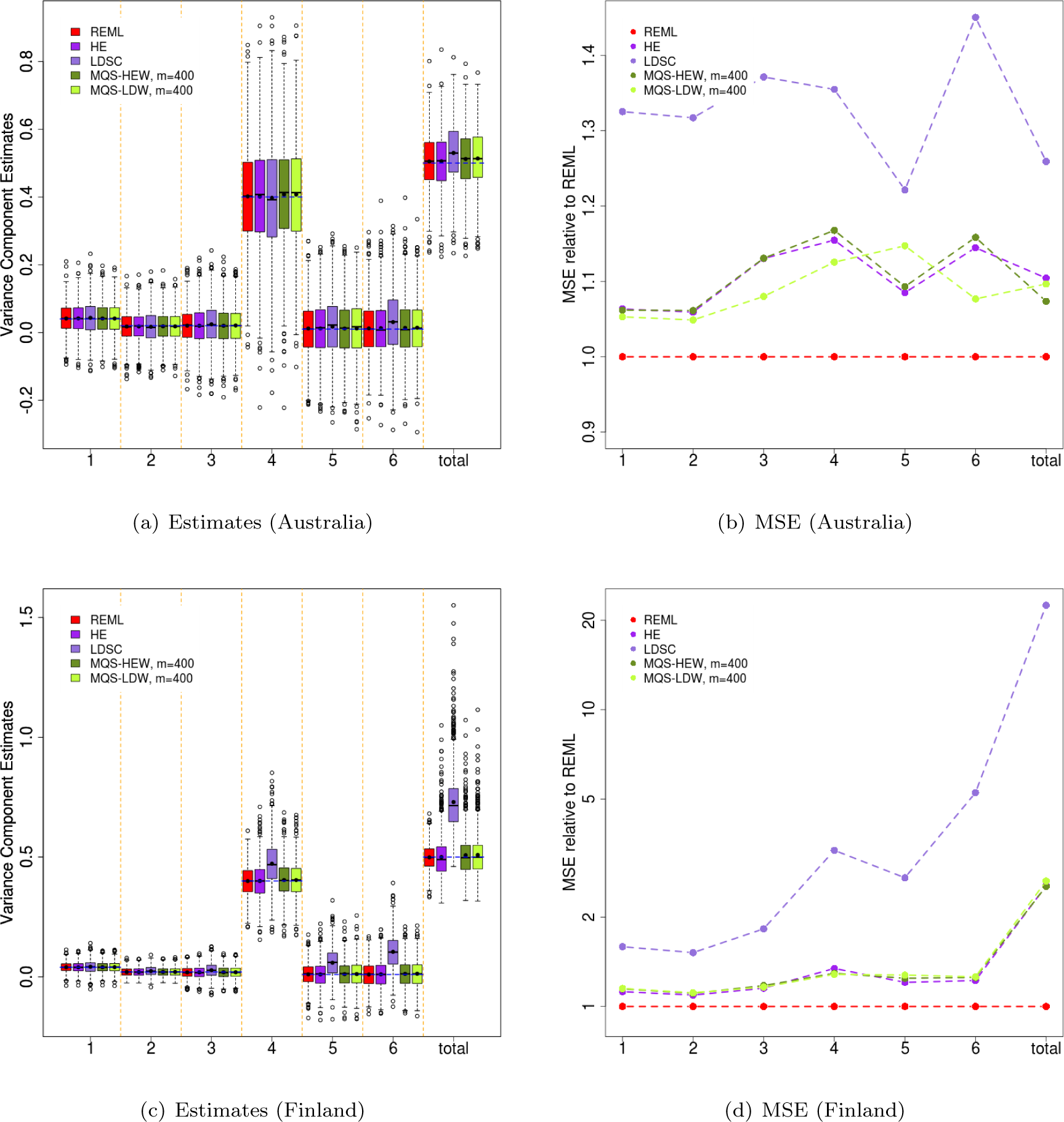
Comparison of variance component estimates from REML (red), HE (purple), LDSC (light purple), MQS-HEW (green), and MQS-LDW (light green) for *k* = 6 simulations based on the Australian data (a, b) or the Finland data (c, d) in scenario III of the *k* = 6 simulations. (a) and (c) show boxplots while (b) and (d) show the MSE relative to REML for all six variance components (1-6) as well as the total. The true variance components are shown as three blue horizontal dashed lines in (a) and (c). The dot in the middle of the boxplots represents the mean of the estimates in (a) and (c). Note that the y-axis in (d) is on log scale.

Finally, we check whether different methods produce calibrated confidence intervals. To do so, we compute the coverage probabilities of the 95% confidence intervals from different methods using the 1,000 replicates from both *k* = 1 simulations and *k* = 6 simulations (Figure 3). If the confidence interval is calibrated, then the coverage probability is expected to be 0.95, with 99% chance in the range of (93.1%, 96.7%). We use CI to compute standard errors for HE and we use CI1 or CI2 to compute standard errors for MQS-HEW and MQS-LDW. The results are as expected: REML, CI for HE, CI1 for both MQS-HEW and MQS-LDW all produced calibrated confidence intervals. CI2 works well in the Australia data with short range LD, but works poorly in the Finland data with long range LD. Like CI2, LDSC produced calibrated confidence intervals in the Australia data but not in the Finland data. The failure of LDSC confidence intervals in the Finland data is in part due to the estimation bias and in part due to the approximate standard error computation method.

**Figure 3.**
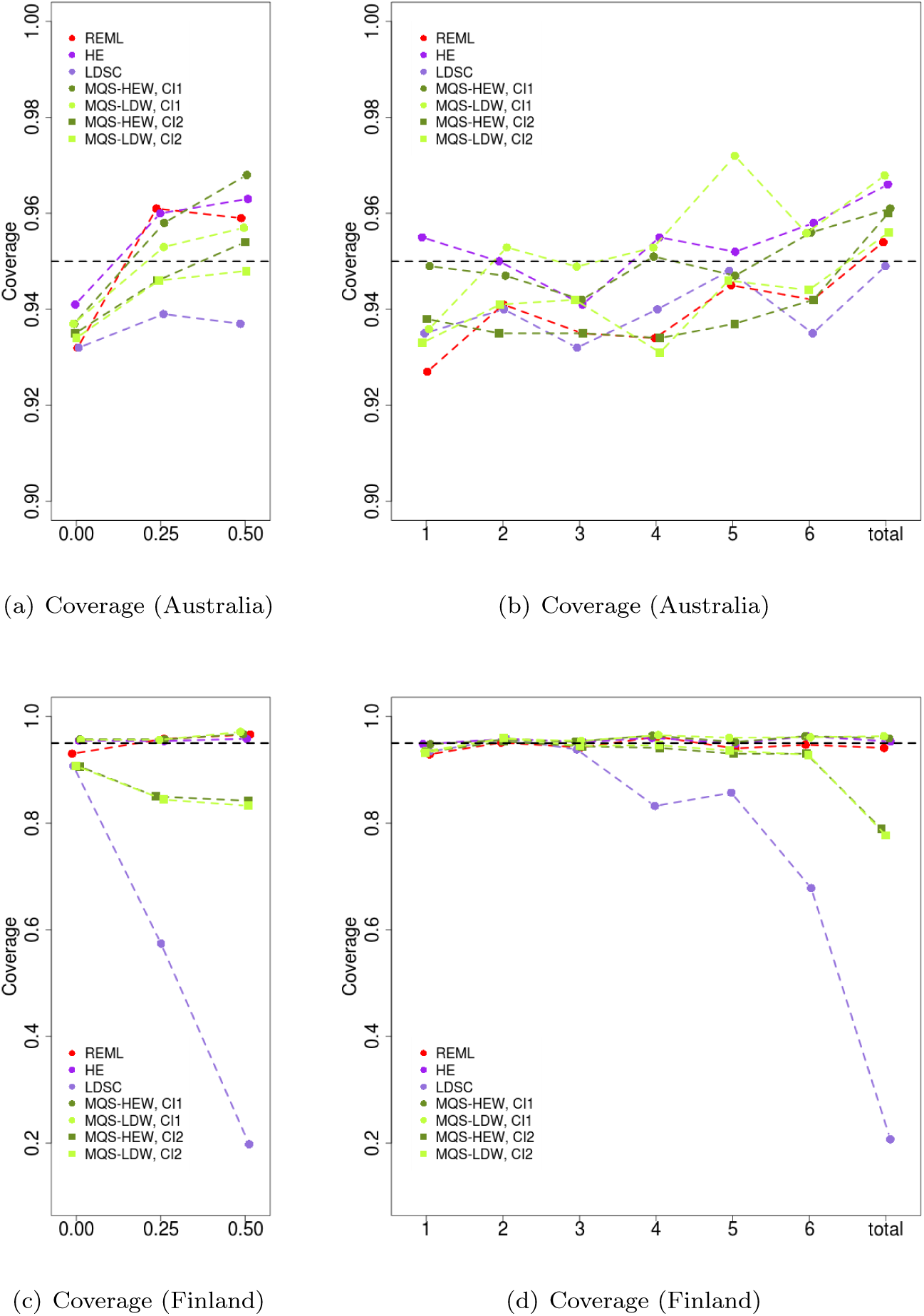
Comparison of the coverage probability of confidence intervals from REML (red), HE with CI (purple), LDSC (light purple), MQS-HEW (green), and MQS-LDW (light green) for simulations based on the Australian data (a, b) or the Finland data (c, d). Two different methods are used for MQS-HEW and MQS-LDW to compute the confidence intervals: CI1 (circle) and CI2 (square). The coverage probability is computed based on a 95% confidence interval. (a) and (c) show the coverage probability for the *k* = 1 simulations, where the true variance components (0, 0.25 and 0.5) are shown on the x-axis. (b) and (d) show the coverage probability for six variance components as well as the total heritability in the *k* = 6 simulations with scenario III. Notice that the scale of y-axis in (a) and (b) is different from that in (c) and (d).

### Real Data Applications

To obtain further insights into the differences between various methods, we apply all five methods to estimate chip heritability for 18 phenotypes in three human GWAS data sets. The first GWAS data is the Australian data that contains height measurements for Australian. The second GWAS data is the Finland data that contains 10 quantitative traits, including C-reactive protein (CRP), glucose, insulin, total cholesterol (TC), high-density lipoprotein (HDL), low-density lipoprotein (LDL), triglycerides (TG), body mass index (BMI), systolic blood pressure (SysBP) and diastolic blood pressure (DiaBP). The third GWAS data is the WTCCC data [54], which includes about 14,000 cases from 7 common diseases and about 3,000 shared controls, all typed on a common set of 458,868 SNPs. The 7 common diseases are bipolar disorder (BD), coronary artery disease (CAD), Crohns disease (CD), hypertension (HT), rheumatoid arthritis (RA), type 1 diabetes (T1D) and type 2 diabetes (T2D). We select the WTCCC data because this is a case control study and we expect long range LD in case control studies due to ascertainment [48, 49]. We apply the five methods to these data in the same way as described in the simulations. For LDSC, we also use 10 MB sliding window in addition to the 1 MB window to estimate the LD scores.

The chip heritability estimates are presented in Table 2. For case control studies, we present estimates on the observed scale, which can be easily converted to the liability scale with a known disease prevalence in the population [21,33,55]. For example, for a disease with a disease prevalence of 0.5% in the population and a case proportion of 50% in the case control study, then the scaling factor is 0.47. Because of the small scaling factor, the heritability estimates for some diseases are above one. The heritability estimates in the Finland data are largely consistent with a previous study [56]. The estimates in the WTCCC are also consistent with a previous study [33]. The heritability estimate for height in the Australia data is slightly smaller than that from previous studies [21,24]. This is because we used imputed data here. Using imputed data is known to disrupt LD pattern and reduce heritability estimates [57].

**Table 2.** Chip heritability estimates from different methods for 11 quantitative traits and 7 binary phenotypes from three GWASs. Values in parentheses are standard errors. For MQS-HEW and MQS-LDW, the standard errors are computed by CI1. The standard errors computed by CI2 are available in a supplementary table. The heritability estimates for the WTCCC data is presented at the observed scale. A small scaling factor is required to transform the estimates to liability scale.

| Trait | REML | HE | Methods | | MQS ($m = 400$; CI1) | |
| --- | --- | --- | --- | --- | --- | --- |
|  |  |  | LDSC |  |  |  |
|  |  |  | 1 MB | 10 MB | HEW | LDW |
| Australia, $n = 3,925$, $p = 4,352,968$ | | | | | | |
| Height | 0.27(0.072) | 0.25(0.072) | 0.30 (0.080) | 0.29 (0.076) | 0.26(0.072) | 0.28(0.072) |
| Finland, $n = 5,123$, $p = 319,148$ | | | | | | |
| CRP | 0.043(0.049) | 0.034(0.045) | 0.054(0.066) | 0.046(0.056) | 0.037(0.045) | 0.037(0.045) |
| Glucose | 0.17(0.046) | 0.21(0.069) | 0.30 (0.067) | 0.25 (0.057) | 0.21(0.070) | 0.21(0.068) |
| Insulin | -0.063(0.037) | -0.084(0.043) | -0.12 (0.057) | -0.1 (0.0476) | -0.082(0.044) | -0.081(0.044) |
| TC | 0.29(0.043) | 0.42(0.097) | 0.63 (0.079) | 0.54 (0.068) | 0.42(0.098) | 0.43(0.099) |
| HDL | 0.34(0.043) | 0.36(0.071) | 0.50 (0.081) | 0.42 (0.067) | 0.36(0.072) | 0.35(0.068) |
| LDL | 0.38(0.041) | 0.60(0.13) | 0.91 (0.080) | 0.77 (0.069) | 0.61(0.13) | 0.61(0.13) |
| TG | 0.19(0.047) | 0.17(0.053) | 0.25 (0.070) | 0.21 (0.059) | 0.18(0.053) | 0.17(0.053) |
| BMI | 0.20(0.047) | 0.18(0.053) | 0.28 (0.063) | 0.24 (0.054) | 0.19(0.053) | 0.19(0.054) |
| SysBP | 0.19(0.045) | 0.27(0.087) | 0.41 (0.064) | 0.35 (0.055) | 0.27(0.088) | 0.28(0.090) |
| DiaBP | 0.071(0.046) | 0.073(0.049) | 0.11 (0.063) | 0.093 (0.053) | 0.075(0.049) | 0.076(0.049) |
| WTCCC, $n \sim 5,000$, $p = 458,868$ | | | | | | |
| BD | 0.83(0.057) | 1.05(0.095) | 1.32(0.085) | 1.31(0.084) | 1.08(0.098) | 1.16(0.10) |
| CAD | 0.57(0.061) | 0.58(0.071) | 0.72(0.078) | 0.71(0.078) | 0.60(0.073) | 0.63(0.074) |
| CD | 0.71(0.060) | 1.06(0.15) | 1.32(0.11) | 1.30(0.10) | 1.08(0.16) | 1.12(0.17) |
| HT | 0.58(0.060) | 0.62(0.072) | 0.78(0.077) | 0.77(0.076) | 0.63(0.072) | 0.66(0.073) |
| RA | 0.69(0.059) | 0.81(0.083) | 0.95(0.32) | 0.90(0.29) | 0.84(0.085) | 0.83(0.083) |
| T1D | 0.97(0.052) | 1.53(0.15) | 1.65(0.74) | 1.48(0.62) | 1.60(0.15) | 1.41(0.11) |
| T2D | 0.61(0.060) | 0.69(0.082) | 0.87(0.080) | 0.86(0.078) | 0.71(0.084) | 0.76(0.084) |

The real data results are consistent with what we see from the simulations. First, MQS-HEW estimates and the standard errors (CI1) are almost identical to that of HE (CI). Second, LDSC estimates are larger than the other estimates, consistent with the upward bias in the simulations. The bias is also more obvious in the Finland and the WTCCC data than in the Australia data. The bias can be reduced by using 10 MB estimation window instead of 1 MB but cannot be completely removed. In contrast, consistent with its known downward bias in case control studies [32,33], REML estimates are consistently smaller in 7 disease phenotypes. Third, MQS-LDW estimates are largely similar to MQS-HEW for quantitative traits and for diseases with a low chip heritability estimate (e.g. CAD, HT, RA, T2D). However, MQS-LDW estimates are noticeably different from MQS-HEW for diseases with a high chip heritability estimate (e.g. BD, CD, T1D). Because we do not know if MQS-LDW provides unbiased estimates in case control studies, we do not know whether such difference was due to a large variance or a potential bias of MQS-LDW in case control studies. Fourth, consistent with simulations, CI2 often produces overly narrow standard errors when compared with CI1 (Table S2). However, for two disease phenotypes (RA and T1D), the standard errors from CI2 are extremely large, suggesting that the calibration issue with CI2 may not always favor one direction. The standard errors of LDSC are similar to that of CI2 but are different from the exact method CI1 for most traits.

Our method requires genotypes from a random sub-sample of the study to estimate *S*. When such a subset of individuals is not available, we can use a reference panel to estimate *S*, so long as individuals in the reference panel can be viewed as a sub-sample of the study. However, a mismatch between the reference panel and the study sample can cause estimation bias. In addition, using a separate reference panel prevents us from using the exact method CI1 to compute the standard errors. Here, we explore the use of genotype data from the 1,000 genomes project [47] for chip heritability estimation in the three GWASs. Specifically, instead of using 400 randomly selected individual from the study sample, we use 503 individuals of European ancestry from the 1,000 genomes project to estimate *S*. The chip heritability estimates for all traits from the three data sets are shown in Table S2. For both the Australia and WTCCC data sets, using the 1,000 genomes data as a reference panel produce similar results. However, for the Finland data set, the estimates from using the 1,000 genomes data are much larger, suggesting a potential over estimation. The results suggest that a match between the reference panel and study sample is critical for accurate estimation. In addition, because we can only use CI2 to compute the standard errors, the standard errors suffer from the same drawback as detailed in the previous paragraph.

Finally, we apply MQS-HEW and MQS-LDW methods to analyze 8 phenotypes from four consortium studies. These phenotypes include BMI (*n* = 120, 569), height (HT, *n* = 129, 945) from the GIANT consortium [58,59], HDL (*n* = 88, 754), LDL (*n* = 84, 685), TC (*n* = 89,005) and TG (*n* = 85, 691) from the Global Lipids Genetics Consortium [60], fasting glucose (FG, *n* = 58,074) from the MAGIC consortium [61], and Crohn’s disease (CD, *n* = 21,447) from the International Inflammatory Bowel Disease Genetics Consortium [62]. The data have been pre-processed by a previous study [63]. We further select a common set of *p* = 5,014, 740 SNPs among these phenotypes for analysis. We partition SNPs into the same six functional categories (coding, UTR, promoter, DHS, intronic and else) as before [29]. Because only z-scores are available for these phenotypes, we have to use CI2 to compute the standard errors and use genotypes from 503 individuals of European ancestry in the 1000 genomes project [47] as a reference panel to estimate **S**. In addition, following previous approaches [30], to contrast the importance of different categories, we focus on estimating the relative value instead of the absolute value of variance components. Specifically, as in [30], we construct a fold enrichment parameter, defined as the ratio between the per-SNP variance in one category and the per-SNP variance in all categories, to quantify the relative importance of different functional categories (Supplementary Text).

Figure 4 shows the enrichment parameters for six categories in 8 phenotypes estimated by either MQS-HEW or MQS-LDW. The results from both MQS-HEW and MQS-LDW are consistent with what we expect [30]: for most phenotypes (with the notable exception of BMI), the per-SNP variance in the coding region is the largest, followed by the UTR, promoter and the DNS regions. The per-SNP variance for both the intronic and intergenic regions are close to zero. The enrichment estimates between MQS-HEW and MQS-LDW are similar overall, though the enrichment of the coding region is estimated to be larger in MQS-LDW than in MQS-HEW for the lipid phenotypes.

**Figure 4.**
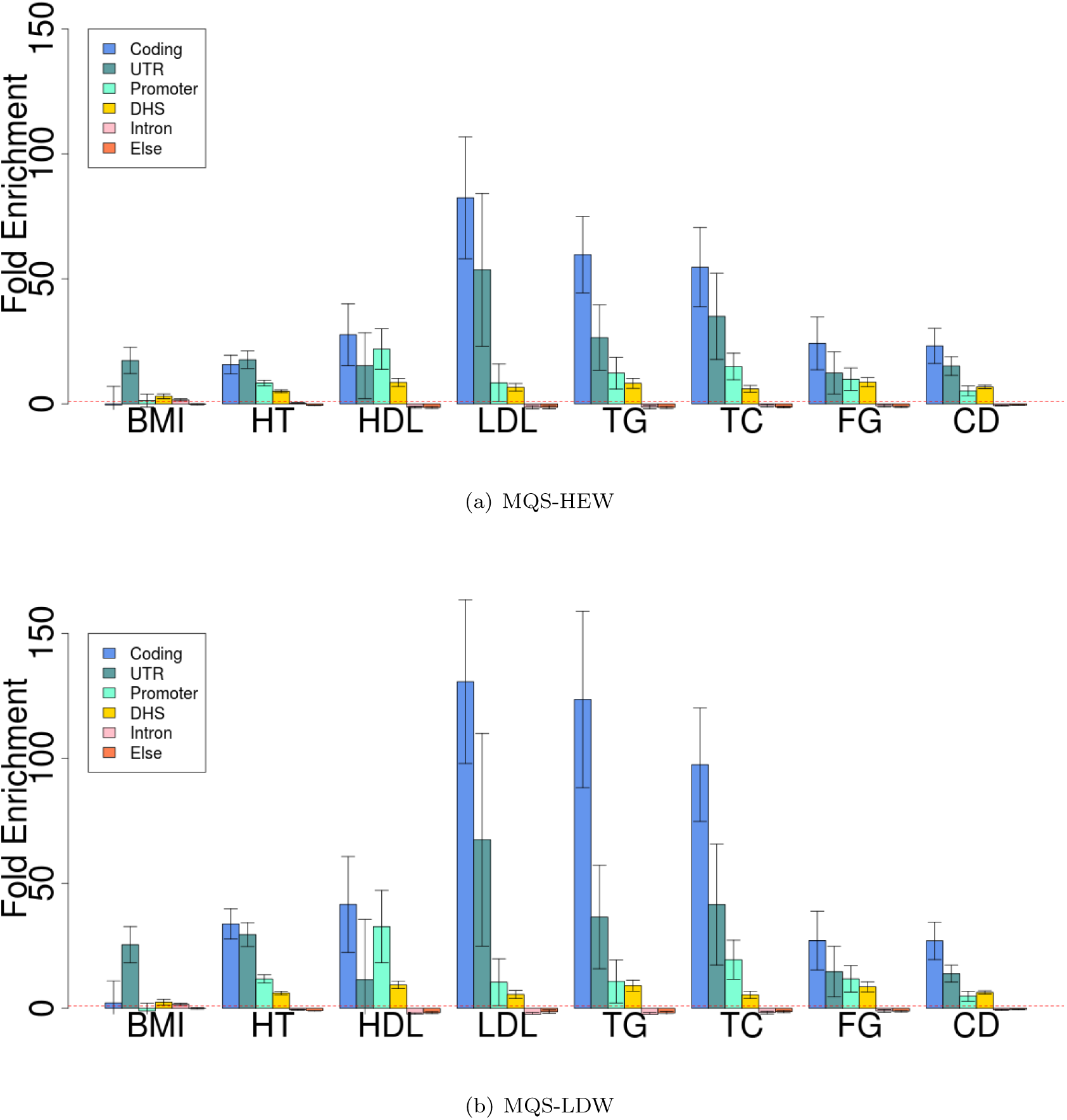
MQS-HEW (a) and MQS-LDW (b) reveal the importance of six functional categories in 8 phenotypes from four GWAS data sets. y-axis shows the fold enrichment, computed as a ratio between the average SNP effect size in one category and the average SNP effect size across the whole genome. Both MQS-HEW and MQS-LDW use marginal z-scores together with genotypes of 503 individuals with European ancestry from the 1,000 genomes project. CI2 is used to construct the confidence intervals.

## Discussion

We have presented a novel framework, MQS, for variance component estimation with summary statistics. MQS is computationally efficient for large data and produces unbiased estimates with calibrated standard errors. MQS is also flexible and can be used with other methods to model uneven linkage disequilibrium [25,64] or model the effect size dependency on minor allele frequencies [21]. In addition, MQS can be extended to model multiple correlated phenotypes [13,46] and/or incorporate overlapping SNP functional annotations. With simulations and applications to 33 phenotypes from 7 GWASs, we have shown the benefits of our method.

We have focused on two variations of MQS in the present study. Both variations, MQS-HEW and MQS-LDW, yield unbiased estimates but with varying degrees of accuracy. Although we cannot tell in advance which method is more accurate for a particular data set, simulations suggest that, when *k* = 1, MQS-LDW may be more accurate than MQS-HEW for quantitative traits and unrelated individuals. The superior accuracy presumably stems from the fact that MQS-LDW uses LD score information and relies on *a priori* set of estimates from MQS-HEW. However, MQS-LDW is not always more accurate than MQS-HEW. In real data applications, the estimates from MQS-LDW are often very similar to that from MQS-HEW. MQS-LDW is also comparable to and sometimes even worse than MQS-HEW for related individuals or for *k* > 1. In addition, MQS-LDW suffers from two important drawbacks. First, because MQS-LDW requires an iterative procedure, computing the standard errors using CI1 for MQS-LDW is less convenient than MQS-HEW. Inconvenience in computing the exact standard errors can affect the usage of MQS-LDW in consortium studies. Second, because of the iterative procedure and the non-linear dependence on the MQS-HEW estimates, it is unclear whether MQS-LDW can produce unbiased estimates in case control studies as MQS-HEW is known to do [32,33]. Therefore, at this stage, we recommend the use of MQS-HEW as a default choice for both quantitative traits and case control studies. However, exploring other variations of MQS by using other SNP weighting matrices, especially non-diagonal ones, will be an interesting avenue for future research. Our derivation of MQS-LDW provides some possible non-diagonal weight matrices for exploration. It would be ideal to identify a weighting matrix that can lead to estimates that are consistently more accurate than MQS-HEW while remaining unbiased in case control studies.

MQS uses a small random subset of individuals to estimate **S**. Using **Ŝ** instead of **S** reduces much of the computational cost while yielding estimates that are almost as accurate as if the full data were used. For instance, in both our simulations and real data applications, we have used ~ 10% of the data to estimate **S**. Using ~ 10% of the data incurs minimal loss of accuracy but results in an effective ~ 100 fold speed gain (because computational complexity scales with *m*^2^). Our sub-sampling approach of using **Ŝ** instead of **S** is motivated by recent genetic studies that make use of a reference panel for genotype imputation [65–68], and more recently, for multi-loci analysis [45,69] (including LDSC [45]): when the full data is not completely observed, these studies rely on a reference panel to impute the missing pieces to construct a complete data. Our approach, however, differs from the previous approaches in two important ways: we actively use a subset of data to estimate certain quantities even when the full data is completely observed (i.e. in line with the idea of stochastic approximation method [70]); and we account for the extra uncertainty introduced by using a smaller subset of data. Importantly, in the present study, we provide an initial set of statistical reasoning to justify our sub-sampling approach in MQS. However, even with our guidelines, it often remains difficult to choose the right number of sub-samples, *m*, for practical analysis. A large *m* would not save much computational time while a small *m* could be insufficient to produce unbiased estimates. In practice, the optimal choice of *m* will likely depend on both the number of categories *k* and the effective number of independent SNPs in each category; thus we caution against the use of an *m* that is too small when *k* is large. Despite this small concern, however, we believe the sub-sampling strategy allows MQS to achieve an appealing balance between computational efficiency and statistical efficiency. With increasing data sizes, exploring the benefits of sub-sampling strategy in other statistical methods for large-scale GWASs – as well as other big data applications – is likely to yield fruitful results in the future.

## Material and Methods

### The Model

Although our method can be reasonably general, we introduce it by considering a particular application: partitioning heritability by different functional categories. To do so, we assume that variants have been pre-classified into *k* different, non-overlapping functional categories. Our goal is to estimate the proportion of phenotypic variance explained (PVE) by all variants in each category. We consider the following model

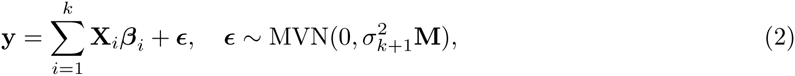

where **y** is an *n*-vector of phenotypes for *n* individuals; **X**_*i*_ is an *n* by *p*_*i*_ genotype matrix for *p*_*i*_ variants in ith category; *β*_*i*_ is a *p*_*i*_-vector of corresponding effect sizes; *ϵ* is a *n*-vector of residual errors; 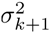 is the residual error variance; 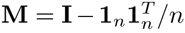 is the projection matrix onto the null space of the intercept; and MVN denotes a (degenerate) multivariate normal distribution. Notice that we ignore the intercept and use **M** instead of the usual identity matrix **I** to reflect the fact that both y and every column of **X** have been centered to have mean zero. Because of the centering, the residual errors follow a multivariate normal distribution with a low-rank covariance matrix **M** that constrains the errors to sum to zero. Centering does not affect results but simplifies the algebra. Generalization of the model to incorporate other covariates in addition to the intercept is straightforward, requiring projecting y, every column of **X** and the residual errors to the null space of the covariates.

Following previous approaches [21,24], for each effect size vector *β*_*i*_, we specify a normal prior with mean zero and variance 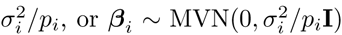. This normality assumption of effect size leads to an alternative but equivalent form – a LMM with *k* variance components [21]

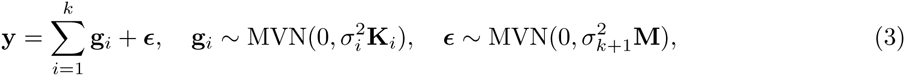

where **g**_*i*_ = **X**_*i*_*β*_*i*_ is the combined genetic effects of *i*th category; 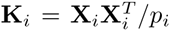 is an *n* by *n* genetic relatedness matrix computed from SNPs in *i*th category; 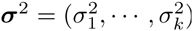 are the variance components.

As usual [24,71,72], we standardize every column of **X** to have variance one. Unlike centering, standardizing the **X** columns will affect the results because it changes our prior assumption [21]. Specifically, standardizing the **X** columns corresponds to making an assumption that rarer variants tend to have larger effects than common variants, and that marker effect sizes depend on the minor allele frequencies (MAFs) in a particular mathematical form (see [21] for relevant discussion). We do not, however, standardize **y**, and the phenotype variance is 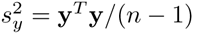.

At this point, it is useful to define the scaled version of the variance components: 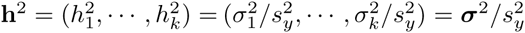 and 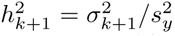. As we will show below, when the phenotype variance 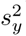 is unknown, ***σ***^2^ and 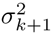 are not estimable from summary statistics alone, but **h**^2^ and 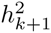 are.

With these modeling assumptions, PVE by ith category is 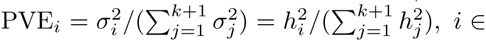, *i* ∈ {1,…, *k*} [21].

Finally, although we will focus on the specific model defined in equation 3, we note that it is straightforward to generalize our model to incorporate other modeling assumptions. For example, we can generalize the model to incorporate other prior assumptions on the dependence of effect sizes on the MAFs (and other variant information), as long as the dependency is linear in 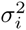 (i.e. in the form of 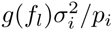, where *f*_*l*_ is the MAF for the *l*th variant in the *i*th category). Such generalization can be achieved by replacing the usual genetic relatedness matrix **K**_*i*_ with a corresponding weighted genetic relatedness matrix with variant weights depending on *g*(*f*_*l*_). Similarly, we can generalize our model to incorporate overlapping categories. If *l*th variant belongs to *k*_*l*_ different categories, we can assume its effect size variance to be a weighted function of the variances of each category, or 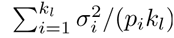. With this assumption, we can again generalize our model by replacing the genetic relatedness matrix **K**_*i*_ with a corresponding weighted genetic relatedness matrix with variance weights 1_*l*∈*i*_/*k*_*l*_, where the indicator function 1_*l*∈*i*_ equals one when the *l*th variant belongs to the *i*th category and equals zero otherwise.

### MoM and MINQUE

Our goal is to estimate the variance components ***σ***^2^ and the residual error variance 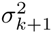 (or the scaled version **h**^2^ and 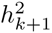). The estimated parameters can then be used to compute PVE. To estimate the variance components, we consider the method of moments, which is based on the following set of quadratic equations (e.g. [34])

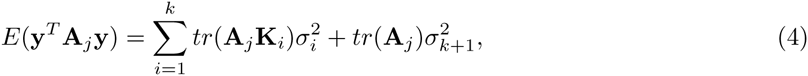

where each **A**_*j*_ is a symmetric non-negative definite matrix and *tr* denotes matrix trace. For estimation with MoM, we replace the expectation on the left hand side (LHS) of equation 4 with the realized value **y**^*T*^ **A**_*j*_**y**. We then solve the equations to obtain estimates for ***σ***^2^ and 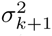. Because there are *k* + 1 parameters in our model, we only need *k* + 1 different **A**_*j*_ to obtain the estimates. Though unnecessarily, we can also use more than *k* +1 **A**_*j*_ to set up an over-determined linear system, and use the ordinary least squares (OLS) to obtain unbiased estimates. For example, the naive LDSC equation [45] is based on using a set of *p* different **A**_*j*_, where **A**_*j*_ = **XΛ**_*j*_**X**^*T*^ and **Λ**_*j*_ is a rank one matrix with *jj*th diagonal element being one and all other elements being zero. (Notice that we call this naive LDSC to distinguish it from LDSC. LDSC does not use OLS estimates based on the naive LDSC equation because of poor performance. Rather, LDSC introduces two extra weights to improve estimation accuracy. See Supplementary Text for details.)

Any choice of **A**_*j*_ will yield unbiased estimates for ***σ***^2^ and 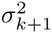, but different choices of **A**_*j*_ affect estimation accuracy. The optimal choice of **A**_*j*_ is found based on the Minimal Norm Quadratic Unbiased Estimation (MINQUE) criterion [34,35,73,74] and takes the following form

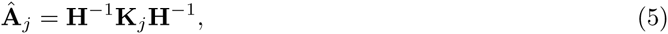

where *j* = 1,…, *k* + 1, **K**_*k*+1_ = **M** and 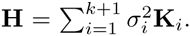. Notice that we have loosely used the matrix inverse notation to denote a generalized inverse, and we have also loosely used hat on top of **A**_*j*_ to not denote an estimate but to denote an optimal choice. Under normality assumptions, MINQUE is also referred to as the best quadratic unbiased estimation (BQUE) [75] and/or the minimal variance quadratic unbiased estimation (MIVQUE) [73,74].

This optimal **Â**_*j*_ depends on a set of variance components that are unknown *a priori.* Thus, we cannot use the optimal **Â**_*j*_ directly in practice. Two options are available to obtain MINQUE estimates in practice. The first option is to apply an iterative procedure on equation 4: in each iteration *t* we plug in the current estimates 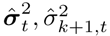 in **H** and **A**_*j*_ to obtained the updated estimates 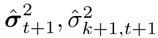. The resulting algorithm is often referred to as an iterative MINQUE, or I-MINQUE, which, not surprisingly, is REML [36]. The second option is to use a pre-determined set of parameters 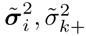 to construct 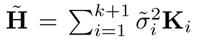 and then use **Ã**_*j*_ = **H̃**^‒1^**K**_*j*_**H̃**^‒1^. For example, MINQUE(0) sets **h**^2^ = **0**, 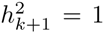 to obtain estimates of reasonably accuracy [76,77]. However, both these two options have the same drawbacks mentioned in the introduction.

### First Approximation: SNP Weights and Approximation to MINQUE

We aim to develop an approximation form of the optimal **Â**_*j*_ that allows the use of summary statistics, facilitates computation, and is reasonably accurate and thus maintains the statistical efficiency comes with the optimal **Â**_*j*_. The particular approximation we consider takes the following form

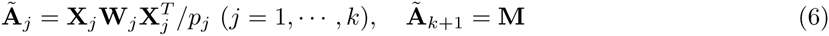

where **W**_*j*_ = *diag*(*w*_*j*1_,…, *w*__*jp*__*j*__) is a pre-specified *p*_*j*_ by *p*_*j*_ diagonal matrix of SNP weights; and **X**_*i*_ again is an *n* by *p*_*i*_ genotype matrix for *p*_*i*_ variants in *i*th category. **Â**_*j*_ (*j* ≠ *k* + 1) is effectively a weighted genetic relatedness matrix, while **Ã**_*k*+1_ approximates **Ã**_*k*+1_ by assuming that **H** is approximately **M**. Notice that **Ã**_*k*+1_ ensures that 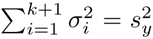 and 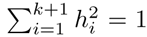. Because a scale transformation of **W**_*j*_ does not affect results, to simplify the algebra, we constrain *tr*(**W**_*j*_) = *p*_*j*_ or equivalently that the average SNP weight equals one.

With this set of **Ã**_*j*_, the set of *k* + 1 estimating equations from equation 4 become

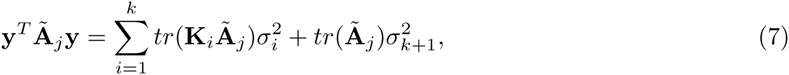

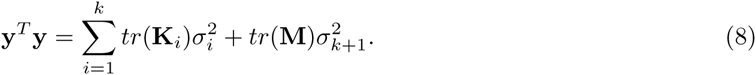

where *tr*(**K**_*i*_) = *tr*(**M**) = *tr*(**Ã**_*j*_) = *n* — 1. We refer to the above equations as MQS estimating equations.

We are yet to specify the weighting matrices **W**_*j*_. In the present study, we consider two particular choices. The two choices represent different ways of approximating the optimal **Â**_*j*_ and are related to the HE regression [32,39–44] and the LDSC [30,45,46], respectively.

### HE Weights

The first set, which we refer to as the HE weights, assigns equal weights for all SNPs, or

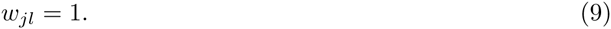

The HW weights are derived based on an approximation to ensure **Ã**_*j*_ ≈ **Â**_*j*_ (Supplementary Text). The approximation becomes exact for unrelated individuals or a trait with zero hertiablity.

We refer to the variation of MQS under the HE weights as MQS-HEW. Surprisingly, MQS-HEW is equivalent to both MINQUE(0) [76,77] and the HE regression (Supplementary Text). Because of the equivalence between MQS-HEW and HE, MQS-HEW is expected to provide unbiased estimates for case control studies [33]. Compared with HE and MINQUE(0), however, our MQS formulation allows the use of summary statistics to compute both the point estimates and the standard errors (see below).

### LD Weights

The second set, which we refer to as the LD weights, takes the following form

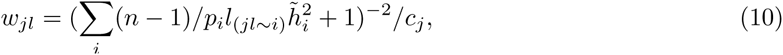

where 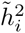 is a pre-specified estimate of 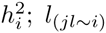 is the LD score of SNP *jl* with respect to all SNPs (including itself if *j* = *i*) in *i*th category; and *c*_*j*_ is a normalizing constant of *j*th category to ensure that Σ_*l*_*w*_*jl*_ = *p*_*j*_. In practice, we use the variance component estimates from MQS-HEW as 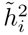, but restricted them to be between 0 and 1 for algorithm stability. The LD score here is defined to be the summation of squared correlations between the *jl*th SNP and all SNPs in the *i*th category minus the expectation under the null, or 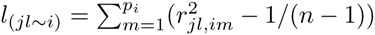. The LD weights are derived based on two approximations to ensure **Ã**_*j*_ ≈ **Ã**_*j*_ (Supplementary Text). The approximations become exact also for unrelated individuals or a trait with zero hertiablity.

We refer to the variation of MQS under the LD weights as MQS-HEW. When *k* = 1 and when the LD scores are exact, MQS-LDW is mathematically equivalent to LDSC [45] (Supplementary Text). However, LD scores in practice are estimated via a sliding window based approach [45] and are inevitably under-estimated. Under-estimation of the LD scores have different impacts on MQS-LDW and LDSC. For MQS-LDW, the estimates with estimated LD scores are still unbiased but are less accurate than that based on exact LD scores. However, because LDSC uses estimated LD scores to approximate *tr*(**K**_*i*_**Ã**_*j*_) instead of computing it as in MQS-LDW, LDSC is expected to under-estimate *tr*(**K**_*i*_**Ã**_*j*_) and thus overestimate the variance components.

When *k* > 1, MQS-LDW is not longer equivalent to the stratified LDSC [30] even with exact LD scores. Unlike *k* = 1, it is no longer possible to convert MQS-LDW estimating equations to a set of linear equations that relate per-SNP z-scores to per-SNP LD scores. Similar to the case of *k* =1, the stratified LDSC estimates are expected to be biased. This time, because the LD scores for SNPs in different categories are inevitably under-estimated to a different degree, the direction and degree of bias in the stratified LDSC are expected to vary across multiple variance components.

### Point Estimates and Confidence Intervals

With the MQS estimating equations 7 and 8, it is straightforward to obtain variance component estimates. To do so, we subtract equation 8 from each equation in 7, solve the resulting *k* equations to estimate **σ**^2^, and plug in the estimated **σ**^2^ to equation 8 to estimate 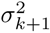. The resulting MQS estimates are in a simple, closed-form solution

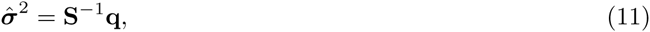

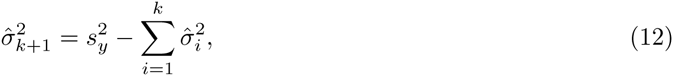

where the elements in the *k*-vector **q** and the *k* by *k* matrix **S** are

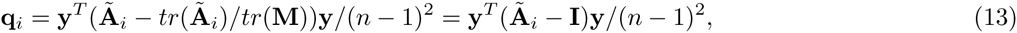

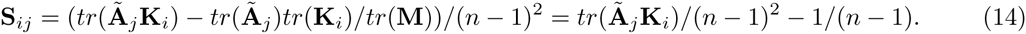

The variance for the estimates are

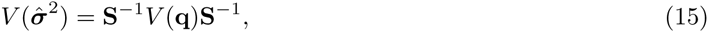

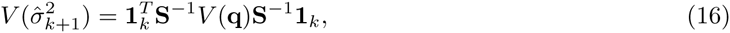

where 1_*k*_ is a *k*-vector of 1s and the *k* by *k* covariance matrix ***V***(**q**) is

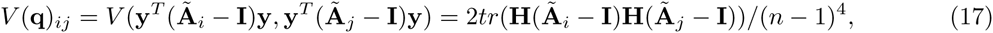

since ***V***(**y**) = **H**.

The above form of ***V***(**q**) requires cubic operations to compute. To speed up computation, we instead consider a novel approximation to ***V***(**q**)

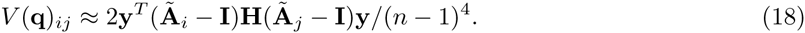

We refer to the above method of computing standard errors as CI. The above approximation is based on replacing the expectation *H* = *E*(**yy**^*T*^) with its realized value **yy**^*T*^. The approximation is motivated by the average information (AI) algorithm [78] and effectively uses a realized information matrix in place of the expected information matrix. Our approximation not only makes computation *n* times faster than the usual MoM, but also allows the use of summary statistics in computing standard errors (see below). We will show later in the simulations that the confidence intervals constructed based on this approximation are indeed calibrated. (As a side note, when the sample size is not large enough and the asymptotic normality does not kick in, then we cannot use ***V***(**q**) for hypothesis tests. Instead, we need to use a mixture of chi-square distributions to obtain more accurate *p*-values [79]. We do not explore this issue here as we typically have large sample size in GWASs.)

### Second Approximation: Estimating S via Sub-sampling

Up to now, the total computational complexity of MQS scales quadratically with respect to the sample size and linearly with respect to the number of SNPs, or on the order of *O*(*pn*^2^). In particular, it takes *O*(*pn*) time to compute **q**, *O*(*pn*^2^) time to compute **S**, and *O*(*n*^2^) time to compute ***V***(**q**). Because the most computationally expensive part is the computation of **S**, we consider estimating **S** instead of computing it. Specifically, we consider using *m* randomly selected individuals from the full sample to estimate **S**, or

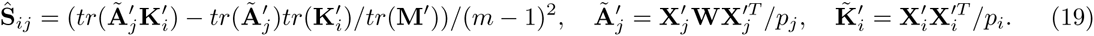

where 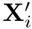 is the standardized genotype matrix for the subset of individuals and **M**′ = **I** – 1_*m*_1_*m*_/*m* is the projecting matrix onto the null space of the intercept for this subset of individuals.

Using the estimated **Ŝ** in place of **S** reduces computational complexity from *O*(*pn*^2^) to *O*(*pm*^2^), resulting in an overall computational complexity of *O*(*pn* + *pm*^2^) for MQS. In addition, using **Ŝ** allows us to apply MQS to data from many consortium studies: we can pair **q** computed from the complete data with **Ŝ** estimated from a random sub-sample of the study. When such a random sub-sample of the study is not available, we can also use a reference panel, such as the 1,000 genomes project [47], to estimate **S**, so long as individuals in the reference panel can be viewed as a sub-sample of the study (e.g. of the same ethnic origin).

To account for the extra variance introduced by using **Ŝ** instead of **S**, we adjust the variance for the estimates by the Delta method

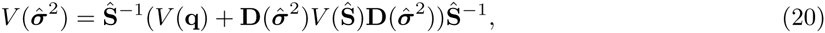

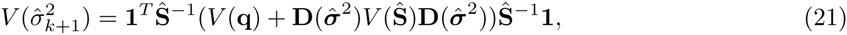

where **D**(**σ̂**^2^) denotes a *k* by *k* diagonal matrix with *i*th diagonal element 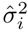. We compute ***V***(**Ŝ**) via Jackknife [80]. Specifically, we first compute 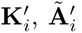 and **Ŝ**. Then, we remove one individual at a time and use the corresponding sub-matrix 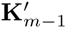 and 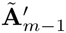 to compute **Ŝ**_*m*–1_. Finally, we estimate the element-wise variance of **Ŝ** as ***V***(**Ŝ**). Because the second step of computing **Ŝ**_*m*–1_ re-uses many quantities that are available from computing **Ŝ**, the overall complexity of estimating ***V***(**Ŝ**) is only *O*(*m*^3^).

Our reasoning behind estimating **S** stems from the following alternative representation of the MQS solution

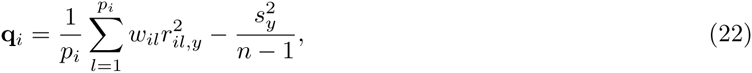

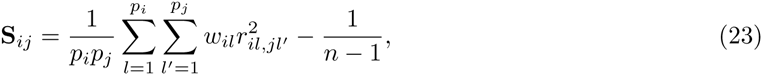

where 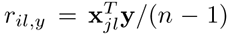 is the correlation between phenotype and the genotype of *l*th SNP in *i*th category, and 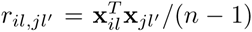 is the genotype correlation between *l*th SNP in *i*th category and *l*′th SNP in *j*th category. Intuitively, each element of **S** is a weighted average of *p*_*i*_*p*_*j*_ different terms of 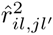 while each element of **q** is a weighted average of only *p*_*i*_ different terms of 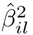. Therefore, **S** is much easier to be estimated accurately than **q**.

Besides the intuitive explanation, we also provide two formal arguments to support the sub-sampling approach for MQS. We use MQS-HEW for illustration. Our first argument is that the variance of **S** estimated by using *m* individuals, or ***V***(**S**), is often small compared with **S** itself. Because of this, the Delta method approximation will be accurate and **Ŝ**^‒1^ can be estimated well by using **S**^‒1^. Specifically, when *k* = 1, for independent SNPs, **S** is expected to be 1/*p* while 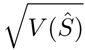 is expected to be 2/(*mp*) under the null, where *p* is the number of SNPs (Supplementary Text). Simulations with independently and binomially distributed SNPs confirm these relationship (Figure S1 and Table S1). For SNPs with LD, we can use the effective number of independent SNPs in place of *p* to provide approximate forms *E*(*S*) ≈ 1 /*p*′ and 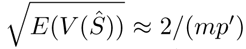. Simulations with real data confirm these approximate relationship (Figures S2, S3 and Table S1).

Our second argument is that ***V***(**Ŝ**) is also small compared with ***V***(**q**). Because of this, the variance of **σ̂**^2^ and 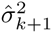 is dominated by ***V***(**q**) and estimating **S** does not introduce much extra variance. Specifically, when *k* = 1, for independent SNPs, 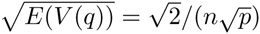 under the null (Supplementary Text), which is 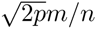 times larger than 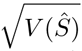. Simulations with independently and binomially distributed SNPs confirm this relationship (Figure S1). For SNPs with LD, we use the effective number of independent SNPs in place of *p* to provide an approximate form 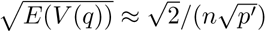. Simulations with real data again confirm the approximate form (Figures S2, S3 and Table S1).

The two arguments above represent the first attempt to understand the behavior of sub-sampling strategy and the reference panel idea that has been widely used in genetics [45,65–69]. However, we note that the two arguments are by no means complete. For example, we have focused on *k* = 1 here. For *k* > 1, the number of variance components *k* as well as the effective number of independent SNPs in each category will both play a role in determining the size of *m*. A larger *m* is likely required to ensure accurate estimation of **S**^‒1^ as well as a small ***V***(**Ŝ**) with respect to ***V***(**q**). In addition, we have focused only on MQS-HEW. MQS-LDW uses an iterative procedure that makes it harder for a thorough investigation. However, we will provide simulations as well as real data applications to show that estimating **S** does work well for either *k* = 1 or *k* > 1 and for both MQS-HEW and MQS-LDW.

### Estimation with Summary Statistics

We are now ready to describe the details of MQS estimation when we only have summary statistics. In this case, we require marginal z-scores from the complete data and individual-level genotypes from a subset of individuals (or from a separate reference panel whose individuals can be thought of as a subset of the study sample). When we only have summary statistics and when the phenotype variance 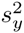 is unknown, **σ**^2^ and 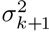 are not estimable but their scaled versions **h**^2^ and 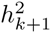 are.

To estimate **h**^2^ and 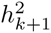, we first use the marginal z-scores to approximate **q**

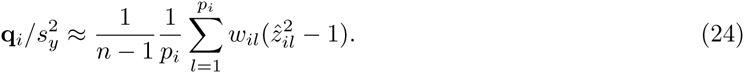

The approximate assumes that the marginal variant effect size is small, which holds well for most GWASs.

Next, we use the individual-level genotype data from a sub-sample of data or a separate reference panel to compute **Ŝ** (equation 19). With 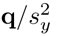 and **Ŝ**, we estimate **h**^2^ and 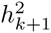 with equations 11 and 12.

To compute the variance for the estimates, we consider two alternative strategies. The first strategy, which we refer to as CI1, is accurate but requires summary statistics in addition to the marginal *z* scores.

This strategy is based on an alternative expression of the equation 18

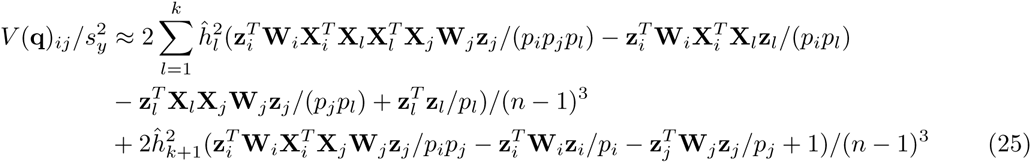

where **z**_*i*_ is a *p*_*i*_ vector of marginal z-scores for SNPs in ith category. Thus, if we can compute additional summary statistics – specifically the *n* by *k* matrix of **X**_*i*_**z**_*i*_, the *n* by *k* matrix of **X**_*i*_**W**_*i*_**z**_*i*_, and the *p* by *k* matrix of **X**^*T*^**X**_*i*_**W**_*i*_**z**_*i*_ – then we can compute the standard errors. These additional summary statistics are easy to compute; in consortium studies, they can be computed within each sub-study and then combined across studies without sharing individual level data. Importantly, for MQS-HEW, we only need to compute two of these three matrices, **X**_*i*__z__*i*_ and **X**^*T*^**X**_*i*__z__*i*_. These two matrices do not require **W**, which is a function of variance components in MQS-LDW. Thus, MQS-HEW can be much more convenient than MQS-LDW. Finally, we note that, although it is tempting to use quantities computed from a subset of individuals to estimate the confidence interval, we find that 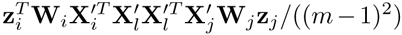 from *m* individuals is not a good estimate of 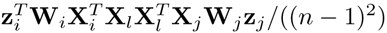.

The second strategy, which we refer to as CI2, does not require extra summary statistics. It is based on the strategy used in LDSC [45]. It works well when SNPs are approximately independent but can work poorly otherwise. This second strategy is based on the observation that the covariance function ***V***(**q**) as a function of **y** can be written as a function of the marginal z-scores **z** = 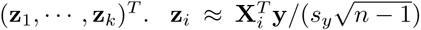 follows approximately a degenerate multivariate normal distribution 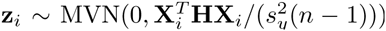. When SNPs are independent (i.e. 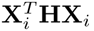 is diagonal) or when 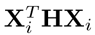 is block-diagonal, we can use block-wise permutation of z-scores to estimate ***V***(**q**) as in LDSC [45]. However, as we show in the Results, when LD pattern is complicated, CI2, like LDSC, can yield untrustworthy confidence intervals.

Overall, we recommend the use of CI1. However, we recognize that CI1 requires additional summary statistics that may not be readily available from many studies at the moment. Thus, we have also implemented CI2 as a useful practical option.

## Acknowledgment

This research is supported by start up funds from the University of Michigan to XZ. We thank Nick Martin and the Queensland Institute of Medical Research for making the Australia height data available to us. We thank Joseph K. Pickrell for making the summary data from 8 quantitative human traits available to us. We thank the NFBC1966 Study Investigators for making the Finland NFBC1966 data available to us. The NFBC1966 study is conducted and supported by the National Heart, Lung, and Blood Institute (NHLBI) in collaboration with the Broad Institute, University of California Los Angeles, University of Oulu, and the National Institute for Health and Welfare in Finland. This manuscript was not prepared in collaboration with investigators of the NFBC1966 study and does not necessarily reflect their views or those of their host institutions. This study also makes use of data generated by the Wellcome Trust Case Control Consortium (WTCCC). A full list of the investigators who contributed to the generation of the data is available from www.wtccc.org.uk. Funding for the WTCCC project was provided by the Wellcome Trust under award 076113 and 085475. We thank Chaolong Wang, William Wen, Ping Zeng, and Xiang Zhu for helpful comments on a previous version of the manuscript.

